# NAT10 Suppresses RNA Sensing Induced IFN-β Transactivation to Promote Viral Infection via Interfering with IRF3 Activities

**DOI:** 10.1101/2025.09.01.673489

**Authors:** Youngmin Park, Jinshan He, Suha Eleya, Zhenyu Wu, Guillaume N Fiches, Dawei Zhou, Emily G Watters, Zhixian M He, Kateepe ASN Shanaka, Thurbu T Lepcha, Yan Liu, Netty G Santoso, Jian Zhu

**Affiliations:** Department of Pathology, College of Medicine, The Ohio State University, Columbus, OH 43210, USA; Department of Microbiology, College of Arts and Sciences, The Ohio State University, Columbus, OH 43210, USA; Department of Microbial Infection and Immunity, College of Medicine, The Ohio State University, Columbus, OH 43210, USA

**Keywords:** NAT10, remodelin, interferon, immune signaling, IRF3, virus, antiviral

## Abstract

Cells can sense invading viruses and trigger type I interferons (IFN-α/β) to evoke antiviral innate immune response. Induction of IFNs needs to be fine-tuned to achieve the antiviral consequence while avoiding severe disruption of host cell homeostasis. Here, we reported that NAT10, the acetyltransferase of histone and N4-acetylcytidine (ac4C) RNA modification, promotes infection of RNA viruses via regulation of type I IFN signaling. Depletion of NAT10 increased the expression of IFN-β and interferon-stimulated genes (ISGs) upon stimulation of type I IFN antiviral signaling, while it impaired viral replication. NAT10 dynamically associated with the IFN-β promotor and negatively regulated IRF3 through modulation of long non-coding RNAs (lncRNAs) that inhibit IRFs. Consistently, the small molecule inhibitor of NAT10, Remodelin, increased IFN-β expression while inhibiting viral infections. Overall, our findings indicated that NAT10 is a negative regulator of type I IFN signaling, suggesting its potential as a target of antiviral treatment.

## Introduction

Type I IFN signaling is an important component of host antiviral innate immunity, which is induced by recognition of pathogen-associated molecular patterns (PAMPs), including viral RNAs. In host cells, viral RNAs are detected by pattern recognition receptors (PRRs), such as Retinoic acid-Inducible Gene I (RIG-I), and this interaction triggers type I IFN signaling. The signal is transduced to downstream adaptor proteins, stimulator of interferon genes (STING) or mitochondrial antiviral signaling protein (MAVS), which trigger its oligomerization and leads to activation of TANK-binding kinase 1 (TBK1). The activated TBK1 directly phosphorylates the IFN regulatory factor IRF3 on multiple Serine/Threonine sties (Ser386/396), which leads to IRF3 dimerization, nuclear translocation to type I IFN promoters, and transactivation of the genes including IFN-β. Secreted IFN-β further binds to receptors on cellular membrane and activates the Janus kinase–signal transducer and activator of transcription (JAK-STAT) pathway, leading to the induction of hundreds of effector ISGs that constitute an overall antiviral intracellular state/environment.

IRF3 is a crucial transcription factor (TF) required for induction of type I IFNs (*1*). Activated IRF3 binds to the IFN-β promoter and leads to its transactivation, which contributes significantly to the antiviral immune responses as well as the maintenance of cellular immune homeostasis (*2*). On the contrary, multiple families of viruses evolve the antagonizing mechanisms that subvert IRF3’s transactivation functions to evade the antiviral immune responses (*3–5*). The association of IRF3 with the IFN-β promoter recruits the histone acetyltransferase/transcription coactivator complex CBP/p300, which subsequently recruits the RNA polymerase II (pol II) and activates transcriptional elongation (*2*, *6*, *7*). Such IRF3-mediated recruitment of RNA pol II involves the chromatin remodeling achieved by CBP/p300-mediated BRG1/BRM-associated factor (BAF) complex of SWI/SNF family, which permits the binding of TATA box-binding protein (TBP) to the promoter of IFN-β (*2*, *7*).

N-acetyltransferase 10 (NAT10), a member of the GCN5-related N-acetyltransferase family, is identified as an acetyltransferase that catalyzes histone and ac4C RNA modifications. Earlier studies showed that NAT10 promotes viral replication of HIV-1, influenza, alpha virus (SINV), EV71, and KSHV (*8–12*) despite that the united underlining mechanisms remain elusive. It is plausible that NAT10 plays a general role in regulating antiviral innate immune responses, given that it has been reported that NAT10 may functionally associate with type 1 IFN signaling in a sepsis mouse model (*13*, *14*). Indeed, more evidence support that NAT10 likely negatively regulates both human innate and adaptive immunity. NAT10 expression is lower in CD4+ T cells of patients with systemic lupus erythematosus (SLE) compared to healthy control (HCs) (*15*). In addition, NAT10 expression is high in multiple types of tumors, indicating that NAT10 may participate in tumor cell-mediated immune evasion in the tumor microenvironment (*16–19*). Thus, we postulated that NAT10 may determine the outcome of viral infections through its functions that regulate immune responses showcased in various pathological conditions.

Our studies identified that NAT10 is a negative regulator of type 1 IFN signaling and directly suppresses the transactivation of IFN-β by interfering with IRF3 through modulation of IRF3-antagonizing lncRNAs, which may cause a general effect to promote viral infections. Overall, our results not only reveal a new role of NAT10 in silencing the antiviral innate immune responses but also shed light in the potential to targeting NAT10 by using its specific small-molecule inhibitors for augmenting antiviral immune responses, which could benefit the treatment of viral infections and associated diseases.

## Results

### NAT10 promotes viral infection through suppression of IFN-stimulated genes (ISGs)

To determine the impact of NAT10 on viral infection, NAT10 was depleted through RNAi-mediated gene knockdown (KD) in the human epithelial cell lines, HeLa-MAGI and A549, which were challenged with influenza (flu) or vesicular stomatitis virus (VSV). The siRNAs targeting NAT10 (siNAT10-1, -2) efficiently depleted endogenous NAT10 protein expression in HeLa-MAGI while resulting in the significant reduction of viral gene expression, measured by protein level for flu NP (**Fig. 1A**) or mRNA level for VSV N transcript (**Fig. 1B**). Likewise, NAT10 depletion also significantly impaired flu infection in A549 cells (**Fig. 1C**). The effect of NAT10 depletion on cell viability of HeLa-MAGI and A549 was negligible as in previous results (**Fig. S1A,B**) (*20*). Importantly, NAT10 depletion also promoted virus-induced expression of ISGs in these cells (**Fig. 1A,C**). We further confirmed the phenotype in the immortalized tracheobronchial epithelial cells (AALE) as well as the primary human small airway epithelial cells (HSAEC) that were transfected with the NAT10 siRNA (siNAT10-2) as well as challenged with flu through measurement of NP and ISG protein expression (**Fig. 1D,E, S1C**). To further validate this, we used the double-color protein immunofluorescence assays (IFAs) to visualize the impact of NAT10 depletion on protein expression of flu NP and ISG15 in HSAEC. The image revealed that the NAT10 KD decreases Flu NP while increasing ISG15 fluorescent signals (**Fig. 1F,G**). Overall, our results showed that the presence of NAT10 promotes viral infection while suppressing induction of ISGs.

**Figure 1.**
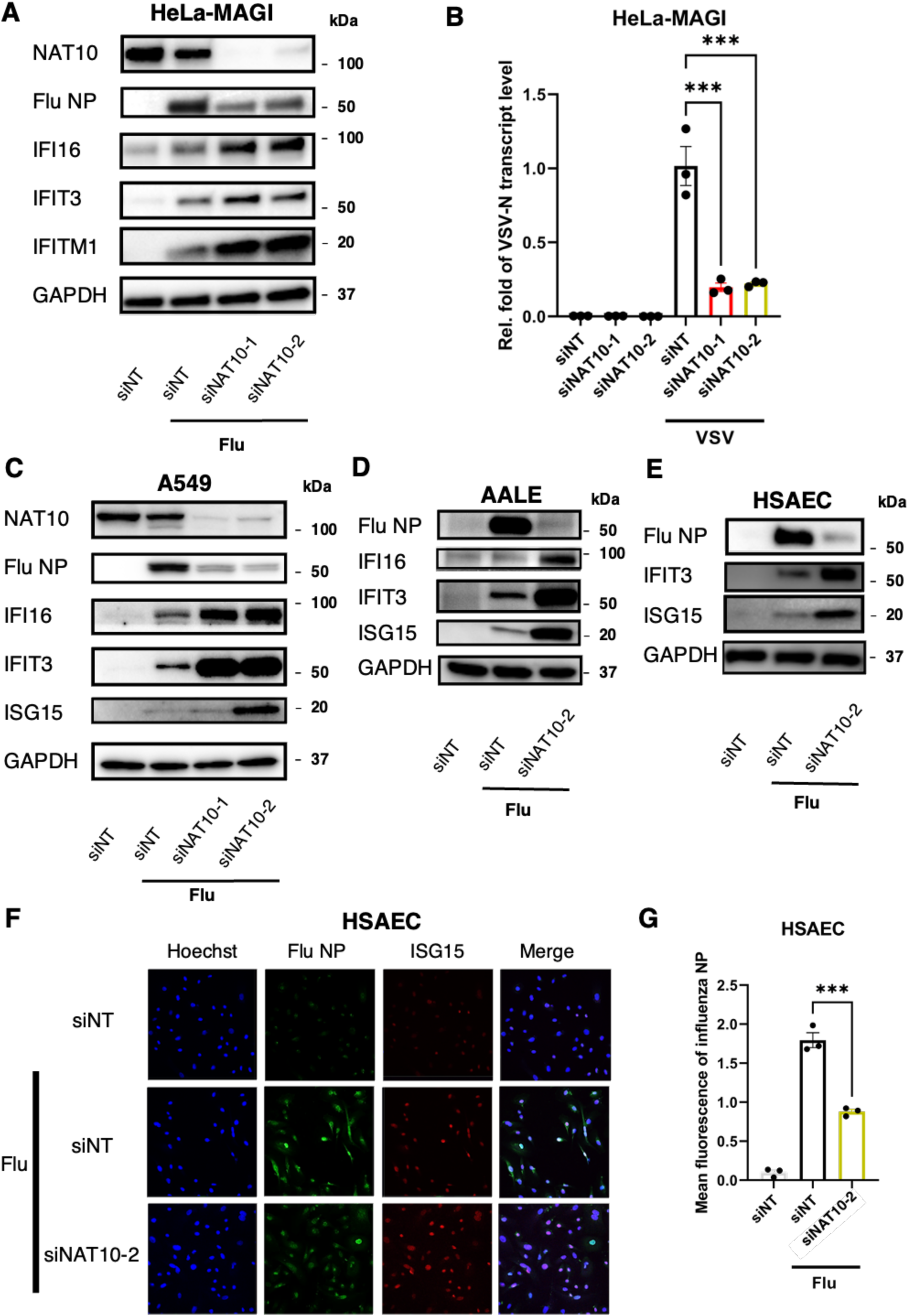
Depletion of NAT10 impaired viral infection while inducing expression of ISGs. (**A**) HeLa-MAGI cells were transfected with NAT10 or non-targeting siRNAs (siNAT10-1, siNAT10-2, siNT), followed by viral challenge with influenza. Expression of viral proteins (flu NP), NAT10, or ISGs were measured by protein immunoblotting using their specific antibodies. GAPDH was used as a loading control. (**B**) HeLa-MAGI cells were transfected with indicated siRNAs, followed by viral challenge with VSV. Expression of viral gene (VSV-N) and NAT10 was measured by RT-qPCR. (**C-E**) A549 (C), AALE (D), or HSAEC (E) cells were transfected with indicated siRNAs, followed by viral challenge with influenza. Expression of viral proteins (flu NP), NAT10, or ISGs was measured by protein immunoblotting. (**F, G**) Protein expression of flu NP and ISG15 was visualized by dual-color immunofluorescence (Alexa 488: flu NP; Alexa 568: ISG15) in A549 cells transfected with the indicated siRNAs and infected with influenza (F). The mean fluorescence intensity (MFI) was quantified by using ImageJ (G). The results were presented as the Mean ± S.E.M of 3 technical repeats from one of at least 3 independent experiment; \*\*\**P* < 0.001, ANOVA and Bonferroni multiple comparison test for (C) or two-tailed Student t-test for (H).

### NAT10 blocks ISG mRNA expression induced by the dsRNA analog poly(I:C)

To further investigate the potential mechanisms for NAT10 to regulate ISG expression, we alternatively stimulated cells with the synthetic double-stranded RNA (dsRNA) analog, the polyinosinic-polycytidylic acid (poly[I:C]) at the optimized doses that activate the viral RNA sensing pathway and induces the antiviral IFN signaling. We noticed that the poly(I:C) treatment has no effect on NAT10 mRNA and protein levels as well as its cellular localization (**Fig. S2A-C**). We showed that NAT10 KD by siRNAs further enhances the poly(I:C)-induced ISG expression in human epithelial cell lines, HeLa-MAGI and A549 (**Fig. 2A,B**), as well as the immortalized and primary human lung epithelial cells, AALE and HSAEC **(Fig. 2C,D**). These results demonstrated that both poly(I:C)- and virus-induced type I IFN responses are consistently regulated by NAT10. We wondered whether NAT10 regulates ISG expression at the transcriptional *vs* translational level, and further quantified the mRNAs for poly(I:C)-induced ISGs by RT-qPCR at the condition of NAT10 depletion. Our results showed that NAT10 KD by its siRNAs significantly decreases NAT10 while increasing mRNA expression of ISGs (IFI16, IFIT3) in poly(I:C)-treated A549 cells **(Fig. 2E-G**). Thus, we concluded that NAT10 blocks viral RNA sensing-induced ISG expression at the transcriptional level.

**Figure 2.**
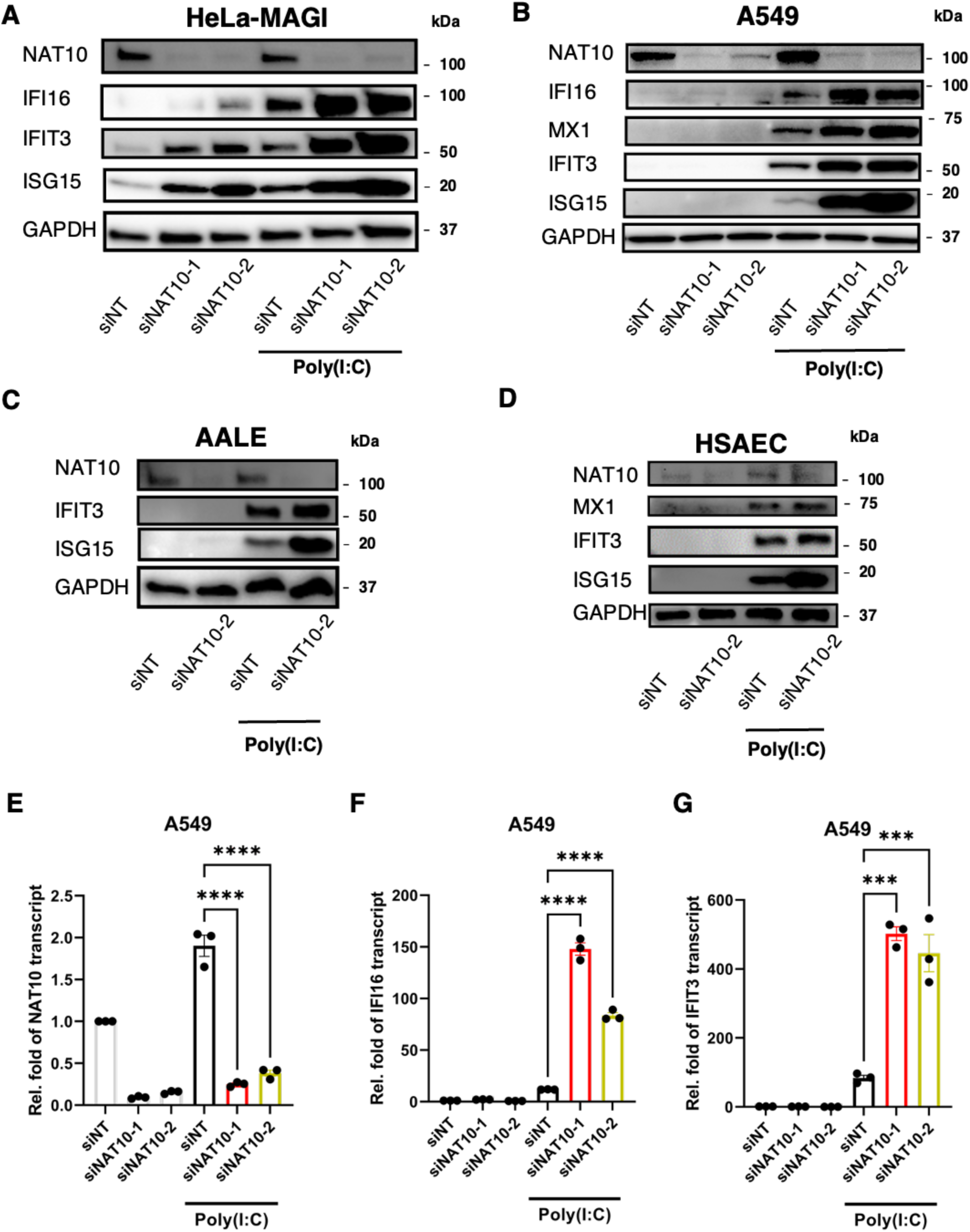
Depletion of NAT10 enhanced the poly(I:C) induced ISG expression. (**A-D**) HeLa-MAGI (A), A549 (B), AALE (C), or HSAEC (D) cells were transfected with NAT10 or non-targeting siRNAs (siNAT10-1, siNAT10-2, siNT), followed by poly(I:C) treatment (1μg/mL). Expression of NAT10 or ISGs was measured by protein immunoblotting using their specific antibodies. GAPDH was used as a loading control. (**E-G**) A549 cells were transfected with indicated siRNAs, followed by poly(I:C) treatment. Expression of NAT10 (E), IFI16 (F), or IFIT3 (G) was measured by RT-qPCR. The results were presented as the Mean ± S.E.M. of 3 technical repeats from one of at least 3 independent experiment; \*\*\**P* < 0.001, \*\*\*\**P* < 0.0001, ANOVA and Bonferroni multiple comparison test for (E-G).

### NAT10 downregulates IFN and ISG expression in type I IFN signaling pathway

To better understand NAT10’s role in controlling antiviral immune responses, we then performed the RNA-seq analysis to characterize the transcriptome subjected to the NAT10-mediated regulation in A549 cells with poly(I:C) stimulation. First of all, our results showed that as expected poly(I:C) treatment induces the upregulation of type I IFNs and ISGs (**Fig. 3A**). IFN-β is considered as a primary type I IFN produced in most non-immune cell types such as epithelial cells (*21–23*). Indeed, IFN-β expression in A549 cells was prominently upregulated due to poly(I:C) treatment compared to IFNα. Enrichment of poly(I:C)-induced type I IFN pathway and other relevant ones were confirmed by Kyoto Encyclopedia of Genes and Genomes (KEGG) and Gene Ontology (GO) analyses (**Fig. S3A,B**). Next, we compared the transcriptomic profiles in poly(I:C)-stimulated A549 cells transfected with either NAT10-targeting or non-targeting siRNAs. It showed that NAT10 KD leads to the remarkable upregulation of IFN-β and ISG transcripts as well as other cytokines/chemokines (**Fig. 3B, C, Table S1**). KEGG and GO analyses revealed that NAT10 depletion upregulates the expression of genes enriched in the type I IFN-related pathways, including cytokine-cytokine receptor interaction, JAK-STAT signaling, toll-like receptor signaling, and RIG-I like receptor signaling (**Fig. 3D, E**). Such enrichment of the type I IFN-related pathways depended on the activation of viral RNA sensing pathway, as it didn’t occur in NAT10 KD cells without poly(I:C) stimulation **(Fig. S3C,D**). Therefore, our transcriptomic analysis identified that NAT10 suppresses the overall type I IFN immune responses triggered by the viral RNA sensing pathway.

**Figure 3.**
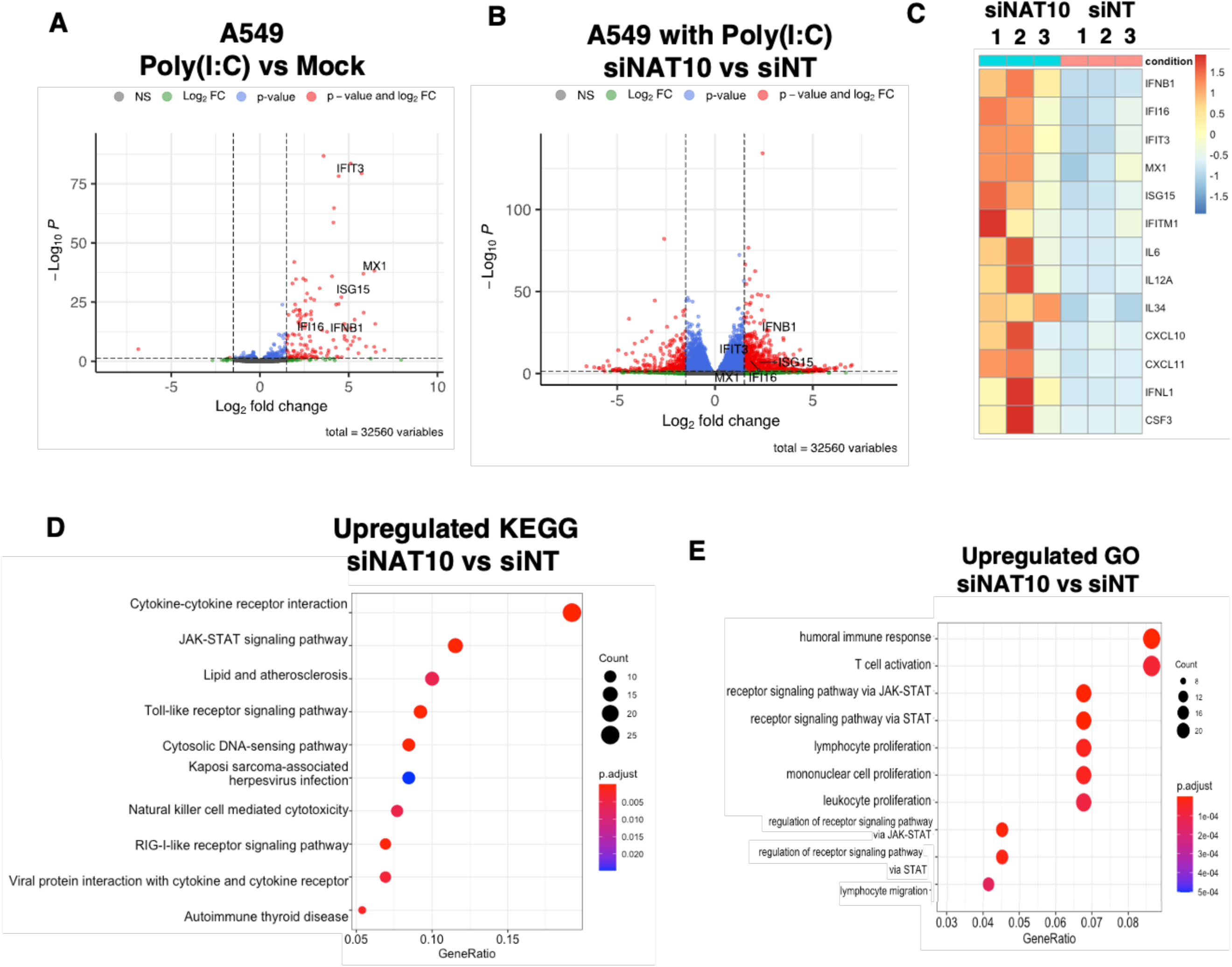
Depletion of NAT10 preferentially induced IFN and ISG expression with poly(I:C) stimulation. (**A-E**) A549 cells were transfected with NAT10 or non-targeting siRNAs (1:1 mixture of siNAT10-1 and siNAT10-2, siNT), followed by poly(I:C) treatment (1μg/mL). Cell samples (triplicate 1-3) were prepared and processed for transcriptomic analysis by RNA-seq. Volcano plot of differentially expressed genes (DEGs) induced by poly(I:C) treatment (A), or NAT10 depletion (B) in A549 cells treated with poly(I:C). Heatmap of the upregulated DEGs due to NAT10 depletion (C). KEGG (D) and GO (E) analysis of top 10 enriched pathway of upregulated DEGs due to NAT10 depletion.

### NAT10 suppresses poly(I:C)-induced IFN-β production in multiple cell types

From the RNA-seq analysis, we noticed that IFN-β is a top hit that is highly upregulated due to NAT10 depletion in poly(I:C)-stimulated A549 cells. It is known that IFN-β is one of the primary cytokines that drive the induction of downstream ISGs through activation of the JAK-STAT pathway (*21–23*). Thus, we postulated that IFN-β is the main gene target of NAT10 to render the suppression of antiviral immune responses. We then confirmed that NAT10 KD by its siRNAs indeed increases the IFN-β production in A549 cells treated with poly(I:C) by both RT-qPCR (**Fig. 4A**) and protein immunoblotting (**Fig. 4B**). We also confirmed that beyond A549 cells NAT10 depletion increases the IFN-β transcript in poly(I:C)-treated HeLa-MAGI cells by RT-qPCR (**Fig. S4A**). We further showed that NAT10 KD by its siRNAs leads to the increase of secreted IFN-β protein in all tested cells stimulated with poly(I:C), including A549 (**Fig. 4C**), HeLa-MAGI (**Fig. S4B**), AALE (**Fig. 4D**), and HSAEC (**Fig. 4E**), by using ELISA. We thus concluded that NAT10 universally suppresses viral RNA sensing-induced IFN-β production in all tested cell types, which would suppress the downstream ISG induction.

**Figure 4.**
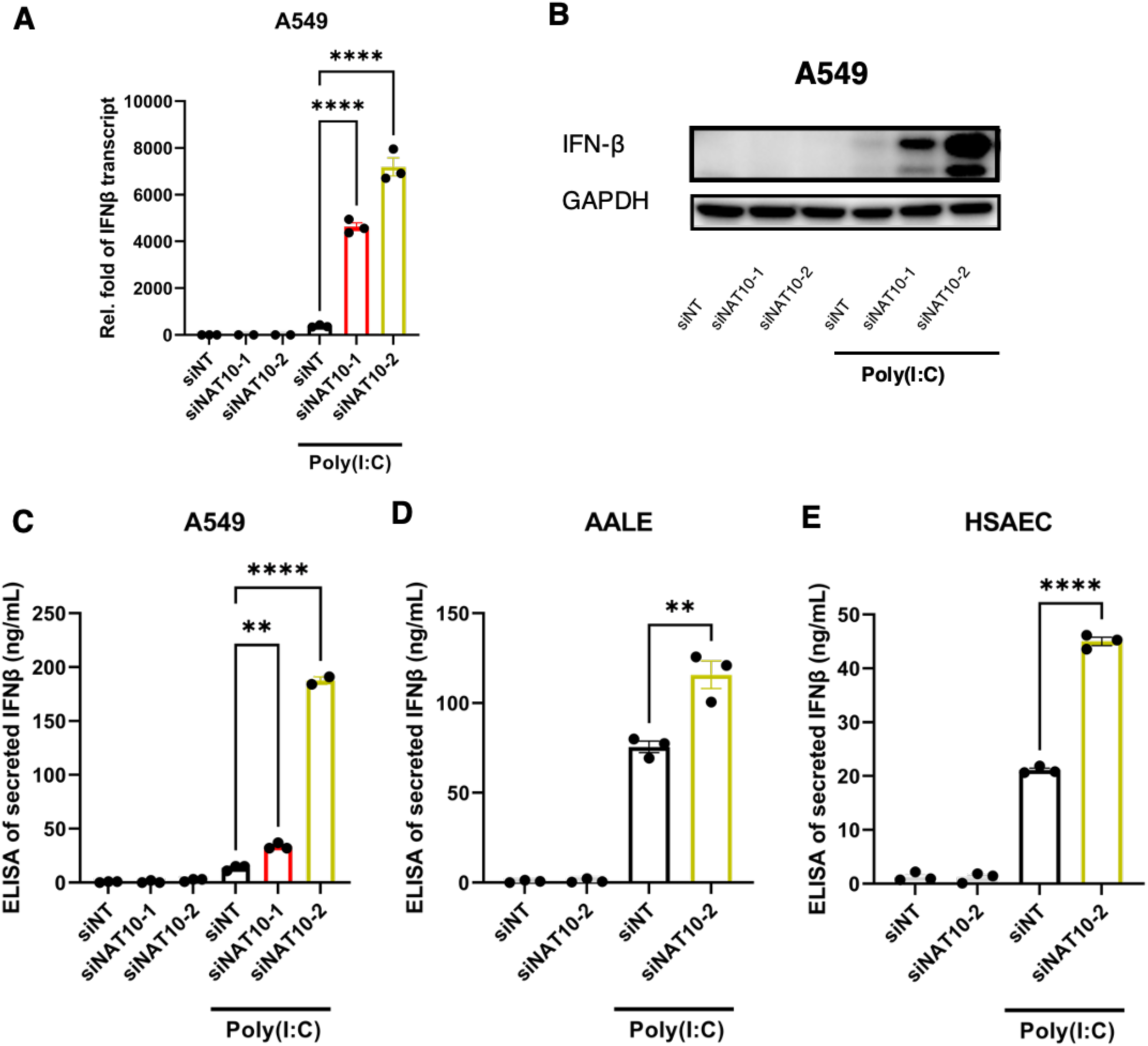
NAT10 suppressed RNA sensing induced transcriptional activation of IFN-β. (**A, B**) A549 cells were transfected with NAT10 or non-targeting siRNAs (siNAT10-1, siNAT10-2, siNT), followed by poly(I:C) treatment (1μg/mL). IFN-β expression was measured by RT-qPCR (A) or protein immunoblotting (B). (**C-E**) Supernatants of A549 (C), AALE (D), or HSAEC (E) cells transfected with indicated siRNAs and treated with poly(I:C) were collected for quantification of secreted IFN-β protein by ELISA. The results were presented as the Mean ± S.E.M. of 3 technical repeats from one of at least 3 independent experiment; \**P* < 0.05, \*\**P* < 0.01, \*\*\**P* < 0.001, \*\*\*\**P* < 0.0001, ANOVA and Bonferroni multiple comparison test for (A, C), Two-way ANOVA for (D, E).

### NAT10 plays a role in silencing IFN-β transactivation via interfering with IRF3

Since our data indicated that NAT10 suppresses IFN-β induction at the transcriptional level, we further performed the luciferase assay by using a vector where the luciferase expression is driven by the IFN-β promoter (*24*). The results revealed that NAT10 KD by its siRNAs increases the relative light units (RLU) of luciferase signal in poly(I:C)-treated HEK293T cells co-transfected with the luciferase/renilla vectors, the RIG-I vector, as well as the siRNAs targeting NAT10 (**Fig. 5A**). Such data indicated that NAT10 imposes an effect on inhibiting the promotor activities of IFN-β. We then decided to identify the underlining mechanisms for NAT10 to silence the IFN-β promotor. We found that NAT10 physically associates with the IFN-β promotor in A549 cells, which is supported by the previously published ChIP-seq data of NAT10 (*25*), and such interaction was significantly reduced due to poly(I:C) stimulation (**Fig. 5B**) or flu infection (**Fig. 5C**).

**Figure 5.**
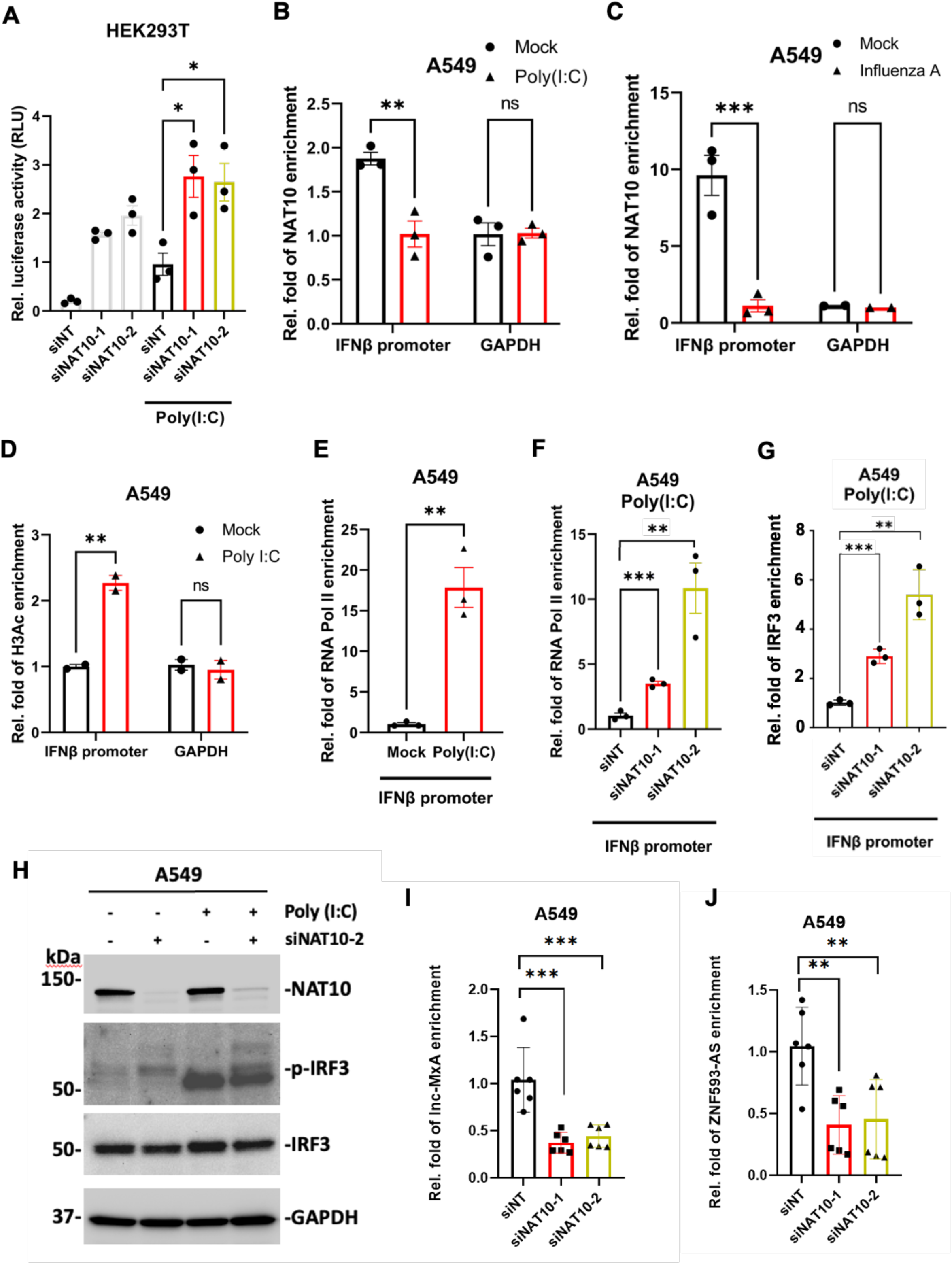
NAT10 contributed to silencing of IFN-β transactivation via interfering with IRF3 activities. (**A**) HEK293T cells were transfected with indicated siRNAs, followed by transfection of IFNβ-luciferase *firefly* reporter vector, *renilla* luciferase vector, and RIG-I vector. Cells were further treated with poly(I:C) (1μg/mL). Luminescence of *firefly* and *renilla* luciferases were quantified, and relative light units (RLU) were calculated. (**B, C**) A549 cells were either treated with poly(I:C) (B) or challenged with influenza (C). Cell samples were prepared and processed for ChIP-PCR to determine NAT10 association at the promoter region of IFN-β. (**D-E)** Cell samples in (B) were processed for ChIP-PCR to determine the H3Ac (D) and RNA Pol-II (E) association at the promoter region of IFN-β. (**F,G**) A549 cells were transfected with indicated siRNAs and treated with poly(I:C) (1μg/mL), which were processed for ChIP-PCR to determine the RNA Pol-II (F) or IRF3 (G) association at the promoter region of IFN-β. (**H**) Total IRF3 and phospho-IRF3(S386) proteins from cell samples in (G) were analyzed by immunoblotting using their specific antibodies. GAPDH was used as a loading control. (**I, J**) A549 cells were transfected with indicated siRNAs, followed by RNA extractions and RT-qPCR assays to quantify the RNA level of lnc-MxA (I) or ZNF593-AS (J). The results were presented as the Mean ± S.E.M. of 3 technical repeats from one of at least 3 independent experiment; \**P* < 0.05, \*\**P* < 0.01, \*\*\**P* < 0.001, ANOVA and Bonferroni multiple comparison test for (A), Two-way ANOVA for (B-E) or two-tailed Student t-test for (F,G).

Notably, NAT10 was initially identified as a histone acetyltransferase that catalyzes histones including the acetylation of histone H3 (H3Ac) that generally associates with the transcriptional activation of genes (*26*, *27*). Thus, we explored NAT10’s role in histone modification near IFN-β promoter and found that NAT10 KD by its siRNAs, while significantly decreasing the intracellular level of total H3Ac, has no obvious effect on the local H3Ac level at the IFN-β promotor nor its chromatin accessibility (**Fig. S5A-C**). However, in contrast to NAT10’s dissociation from IFNβ promoter (**Fig. 5B**), H3Ac level at the IFN-β promotor increased in poly(I:C)-treated A549 cells (**Fig. 5D**), indicating that NAT10 may not be the dominant regulator of histone acetylation at the IFN-β promoter. Interestingly NAT10 depletion further enhanced the recruitment of RNA Polymerase II (Pol II) at the IFN-β promoter induced by poly(I:C) (**Fig. 5E,F**).

It is known that IRF3 is a key transcription factor responsible for the IFN-β induction. IRF3 can be activated due to its phosphorylation by the upstream kinase TBK1. Such activation further leads to IRF3 dimerization and nuclear translocation, as well as its binding to the IFN-β promoter to transactivate its expression by further recruiting RNA Pol II. We thus determined the impact of NAT10 on IRF3 activities. Similar to RNA Pol II, NAT10 depletion by its siRNAs increased the association of IRF3 with the IFN-β promoter (**Fig. 5G**). Depletion of NAT10 also increased the protein level of phosphorylated IRF3 but not total IRF3 in A549 cells in the absence or presence of poly(I:C) (**Fig. 5H**). Correlated with these findings, we observed that depletion of NAT10 also significantly reduced the level of IRF3-antagonzing lncRNAs, lnc-MxA and ZNF593-AS, in A549 cells (**Fig. 5I, J**). Lnc-MxA has been reported to associate with the IFN-β promoter and suppress its transcriptional activation by inhibiting the binding of IRF3 (*5*). It has also been shown that ZNF593-AS directly interacts with the functional domain of IRF3 and suppresses its phosphorylation and activation (*28*). To summarize, our findings unraveled a previous unappreciated mechanism that NAT10 suppresses IFN-β expression by silencing its promotor activities rather through its interference with IRF3-mediated RNA pol II recruitment than its impact on histone acetylation via the acetyltransferase functions.

### NAT10 inhibition induces IFN-β expression while inhibiting viral infection

As we identified that NAT10 promotes viral infection through suppression of IFN-β expression, NAT10 could be considered as a new host gene target for development of antiviral immunotherapies. It has been reported that remodelin hydrobromide, 4-[2-(2-cyclopentylidenehydrazinyl)-4-thiazolyl]benzonitrile, acts as a small-molecule inhibitor of NAT10 by targeting its acetyl group binding site (*29*, *30*). We were intrigued to evaluate the antiviral potential of remodelin in cell cultures. A549 cells were simulated with poly(I:C) and treated with remodelin at a series of doses, which caused the increase of secreted IFN-β measured by ELISA (**Fig. 6A**) as well as the induction of ISGs measured by protein immunoblotting (**Fig. 6B**). We next determined whether the use of remodelin can inhibit the infection of RNA viruses. Our results showed that Remodelin potently blocked VSV infection in A549 cells through measurement of VSV-N expression by RT-qPCR (**Fig. 6C**).

**Figure 6.**
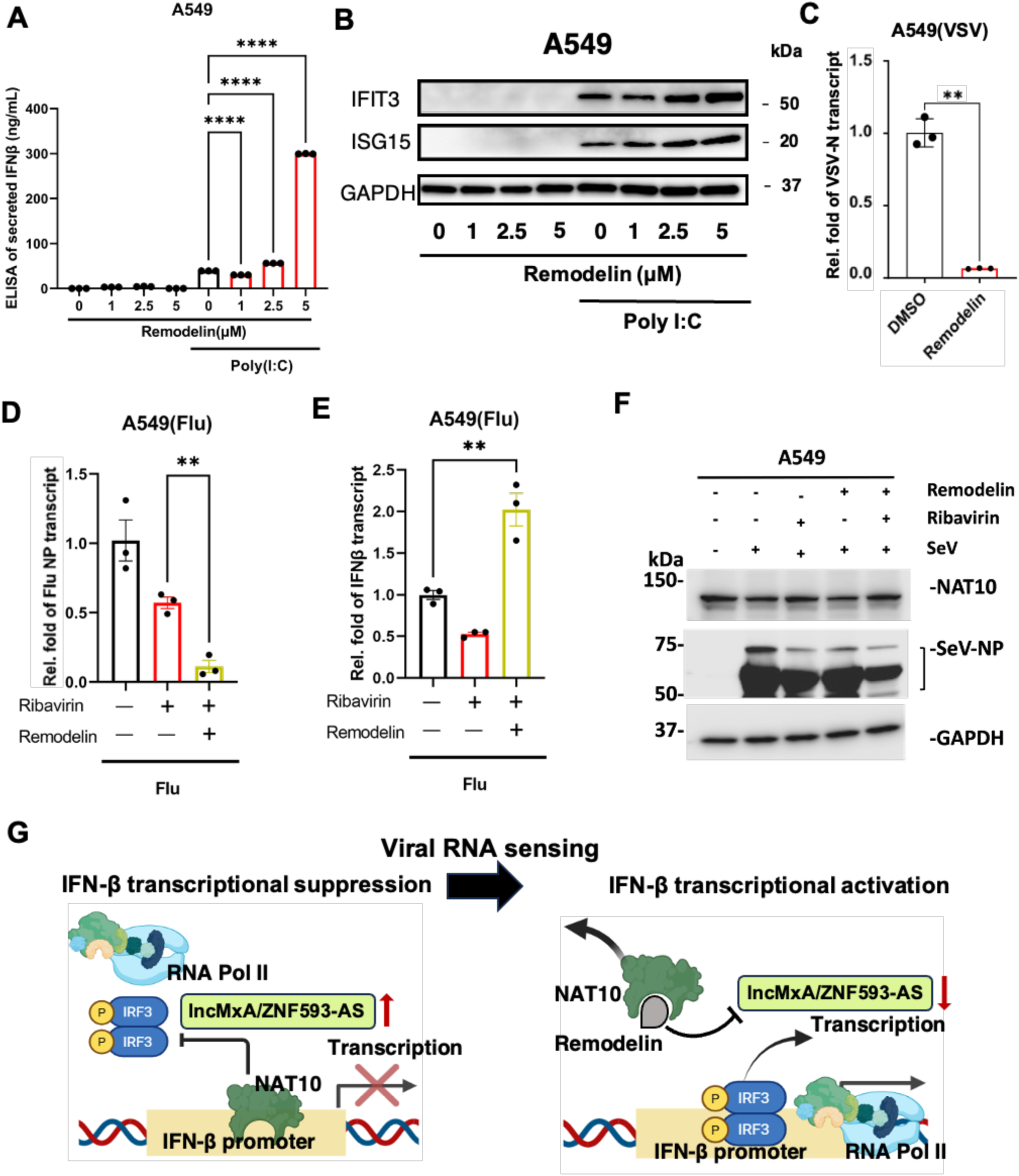
NAT10 inhibitor remodelin induced IFN-β expression while restricting viral infection. (**A**) A549 cells were treated with remodelin at a series of doses (0, 1, 2.5, 5 μM), followed by poly(I:C) stimulation. Cell supernatants were collected for quantification of secreted IFN-β protein by ELISA. (**B**) Cell samples in (A) were subjected to immunoblotting to quantify ISG proteins. GAPDH was used as a loading control. (**C**) Expression of VSV-N in A549 cells infected with VSV and treated with remodelin (5μM) was measured by RT-qPCR. (**D, E**) A549 cells were treated with ribavirin (10μM) alone or in combination with remodelin (5μM), followed by challenge with influenza viruses. Expression of flu NP (D) or IFN-β (E) was measured by RT-qPCR. (**F**) A549 cells were treated with ribavirin (10μM) alone or in combination with remodelin (5μM), and infected with Sendai virus (SeV). NAT10 or SeV-NP protein was measured by immunoblotting using their specific antibodies. GAPDH was used as a loading control. (**G**) A working model that NAT10 suppresses viral RNA sensing induced IFN-β transcriptional activation through interfering with IRF3 activities at the IFN-β promoter by modulating IRF3-antagonizing lncRNAs (lncMxA/ZNF593-AS), which can be disrupted by the NAT10 inhibitor remodelin to argument the anti-viral immune responses. The results were presented as the Mean ± S.E.M.; ns: not significant, \**P* < 0.05, \*\**P* < 0.01, \*\*\**P* < 0.001, ANOVA and Bonferroni multiple comparison test for (A) or two-tailed Student t-test for (C-G).

We also tested the antiviral potency of remodelin on other RNA viruses, including flu and Sendai virus (SeV). However, remodelin treatment alone had no obvious effect on inhibiting flu infection nor induction of IFN-β in remodelin-treated, flu-infected A549 cells (**Fig. S6A,B**). We further tested the anti-flu potential of remodelin in another scenario by combining it with other antiviral reagents, such as ribavirin. Ribavirin is an FDA-approved drug that is used together with pegylated IFNs in clinic as the standard treatment for chronic hepatitis C virus (HCV) infection, which also potently inhibits flu infection (*31*, *32*). We showed that ribavirin in combination with remodelin renders the stronger antiviral potency to block flu NP expression in A549 cells compared to ribavirin alone (**Fig. 6D**), which also restored the drug effect of remodelin to induce IFN-β (**Fig. 6E**). Similarly, the combination of remodelin with ribavirin decreased SeV-NP protein level while neither remodelin nor ribavirin alone had any obvious effect (**Fig. 6F**). Thus, by using remodelin as a chemical probe of NAT10 our studies illustrated that it is promising to target NAT10 for development of pan-antiviral regiments as its inhibition can efficiently induce the activation of antiviral type I IFN signaling.

## Discussion

Recently, there is mounting evidence, including our own data (**Fig. 1-4**), suggesting that NAT10 may play a critical role in regulating virus-host interactions as well as antiviral innate immune sensing and responses (*8–12*). Overall, our studies identified that NAT10 suppresses transcriptional activation of IFN-β that is induced by viral RNA sensing signaling pathway through interfering with IRF3 activities by modulating IRF3-antagonizing lncRNAs, thus favoring infection of RNA viruses (**Fig. 6G**). NAT10 was initially identified as an enzyme that acetylates histones (H3 and H4) and non-histone proteins, but it has also been lately illustrated to catalyze the formation of ac4C at mRNAs (*20*, *27*, *33*). Histone acetylation at multiple lysine sites are generally associated with transcriptional activation (*26*, *34*, *35*), indicating that NAT10’s function to suppress IFN-β might be independent of its histone acetyltransferase activities. Although NAT10 depletion led to the global reduction of total histone H3 acetylation, it appeared to have no effect on reducing local histone H3 acetylation nor chromatin accessibility at the IFN-β promoter (**Fig. S5A-C**). However, H3 acetylation at IFN-β promoter still increased upon poly(I:C) treatment, consistent with previously reported results (**Fig. 5D**) (*7*, *36*).

Our own results showed that NAT10 associates with the IFN-β promoter and that NAT10 depletion enhances the recruitment of RNA Pol II to the IFN-β promoter reminiscent of the similar effect of poly(I:C) stimulation (**Fig. 5B-F**). Thus, we started to explore other potential mechanisms that might involve NAT10-mediated type I IFN suppression. Beyond histones and chromatin remodeling factors, the transcription factor IRF3 can mediate RNA Pol II association and determine the outcome of IFN-β expression. Once IRF3 is phosphorylated and activated, it trans-locates to the nuclei and binds to the IFN-β promoter, which is critical to type I IFN immune responses. We identified an interesting and novel mechanism that NAT10 not only suppresses IRF3 phosphorylation but also reduces its association with the IFN-β promoter (**Fig. 5G,H**). It is plausible that NAT10 acts through its RNA acetyl-transferase functions by targeting RNA species that antagonize IRF3-mediated IFN-β induction. RNA ac4C acetylation by NAT10 may thus play a role in regulating the type I IFN signaling pathway. The evidence has emerged that RNA modifications, including acetylation, contribute to the balance of immune/inflammatory responses as well as the progression of autoimmune and infectious diseases (*37*). It is known that the noncoding RNAs (ncRNAs), including long ncRNAs (lncRNAs), undergo epi-transcriptomic modifications significantly, including RNA acetylation (*38–40*). It is intriguing to characterize NAT10-mediated ac4C modifications of lncRNAs that act as the repressors to antagonize IRF3-mediated IFN-β induction. Indeed, we demonstrated that NAT10 modulates the RNA level of reported IRF3-antagonizing lncRNAs (**Fig. 5I,J**).

Beyond the mechanisms identified from our studies that NAT10 inhibits the IRF3-mediated IFN-β transactivation, there could be alternative mechanisms that potentially explain NAT10’s functions in type I IFN signaling pathway (**Fig. 3**). For instance, it has been reported that NAT10 activates the transcription repressor MORC2 to downregulate CDK1 and Cyclin B1 gene expression during DNA damage (*41*). It has also been shown that p53 is acetylated by NAT10, which impacts p53’s transcriptional activities (*42–44*). There is the evidence that p53 influences the type I IFN signaling by regulating IFN-β expression (*45*). Beyond NAT10’s direct association with the IFN-β promoter to regulate its expression (**Fig. 5**), NAT10 might also interfere with the upstream factors/players in the type I IFN signaling, thus adding another layer of regulatory mechanism. For example, it has been reported that NAT10 dysregulates the STING activation and thus inhibits the type I IFN signaling in a sepsis mouse model (*13*). Therefore, NAT10 likely suppresses IFN-β induction in the type I IFN signaling through multiple mechanisms either directly or indirectly, which requires more detailed investigations.

Since we showed that NAT10 depletion induces the type I IFN signaling while restricting infection of multiple RNA viruses (**Fig. 1, 2**), we speculated that its inhibition would generate the anti-viral benefits. Remodelin is a reported small-molecule inhibitor that targets NAT10’s acetyl-binding site (*46*). We demonstrated that remodelin is capable of not only inducing IFN-β expression **(Fig. 6A,B,E**) but also inhibiting viral infections (**Fig. 6C,D,F**), indicating that its acetyltransferase function could be critical to immune responses and/or viral infections. However, the exact mechanisms of NAT10 inhibition remain not well characterized. Remodelin has been reported to inhibit both NAT10’s RNA and protein acetyltransferase activities (*29*, *41*, *47*, *48*). Detailed biochemical and structural characterizations of NAT10-remodelin interactions would be needed to not only improve the understanding how NAT10 intrinsically regulates its RNA *vs* protein acertyltransferase activities but also optimize remodelin’s chemical scaffold and properties to be further developed as a potential anti-viral drug. We noticed that remodelin is sufficient to inhibit infection of certain RNA viruses (VSV) by its own while it exerts the antiviral effect on others (flu, SeV) only in the combination with ribavirin. It is possible that some of these viruses may employ certain mechanism(s) to antagonize remodelin’s drug effect. It has been reported that the viral proteins of flu, such as PB1, interact with NAT10 (*10*, *49*), which may affect its cellular localization (*50*, *51*). Ribavirin has been implicated to cause the mutagenesis of flu’s viral genome (*52*), which may abolish the viral counteraction against remodelin’s inhibition of NAT10. However, such mechanisms need further studies.

Overall, our study revealed the novel functions of NAT10 to silence the viral RNA sensing-induced antiviral innate immune responses, thus favoring infection of RNA viruses. We further identified a new mechanism for NAT10 to suppress IFN-β transactivation through interfering with IRF3 activities by modulating IRF3-antagonizing lncRNAs. At last, as a proof of principle we showcased that the inhibition of NAT10 by using a small-molecule compound remondlin results in a potent antiviral effect by inducing type I IFN signaling. These findings guarantee the further investigations of NAT10 as a promising host target for augmenting antiviral innate immune responses and the further developments of its small-molecule inhibitors for antiviral immunotherapies.

## Materials and Methods

### Cell culture

A549 (Cat. # CCL-185, ATCC) is cultured in Ham’s F12K (Cat. # 21127022, Gibco). HeLa-MAGI (NIH AIDS Reagent Repository) and HEK293T (Cat. # CRL-3216, ATCC) are cultured in Dulbecco’s modified Eagle’s medium (DMEM, Cat # D5796, Sigma). The cell culture media above contained 10% fetal bovine serum (FBS, Cat. # 10437028, Thermo Fisher), penicillin (100 U/ml) /streptomycin (100 μg/ml) (Cat. # MT30002CI, Corning). HSAEC (Cat. # PCS-301-010, ATCC) are cultured in Airway Epithelial Cell Basal Medium (Cat. # PCS-300-030, ATCC) supplemented with Bronchial Epithelial Growth Kit (Cat. # PCS-300-040, ATCC). Immortalized tracheobronchial epithelial (AALE) cells are cultured in SAGM medium containing supplement kit (Cat. # CC-3118, Lonza).

### Compounds and antibodies

Immuno-stimulant Poly(I:C) (Cat # 528906, Merck) was resuspended in water. Remodelin (Cat # HY16706A, MedChemExpress) was resuspended in DMSO. Ribavirin (Cat. # R0077, Tokyo Chemical Industry) was resuspended in water. The following antibodies were used for protein immunoblotting: anti-NAT10 (Cat # 13365-1-AP, Proteintech), anti-IFI16 (Cat # sc-8023, Santa Cruz Biotechnology), anti-IFIT3 (Cat # sc-393512, Santa Cruz Biotechnology), anti-ISG15 (Cat # sc-166755, Santa Cruz Biotechnology), anti-MX1 (Cat # sc-271399, Santa Cruz Biotechnology), anti-IFITM1 (Cat # 60074-1-lg, Proteintech), anti-IRF3 (Cat # 4302S, Cell Signaling), anti-Phospho-IRF3(Ser386) (Cat # 37829S, Cell Signaling), anti-RNA Polymerase II (Cat. # 39097, Active Motif), anti-GAPDH (Cat # sc-47724, Santa Cruz Biotechnology), anti-H3 (Cat # 61800, Active Motif), anti-H3ac (Cat # 61937, Active Motif), anti-N4-acetylcytidine antibody (Cat # EPRNCI-184-128, ABCAM), anti-NP (Cat # GTX125989, GeneTex), goat HRP-conjugated anti-mouse IgG antibody (Cat. # 7076S, CST), and goat HRP-conjugated anti-rabbit IgG antibody (Cat. # 7074S, CST).

### Virus infection

Sendai virus (SeV) and Influenza A virus (A/WSN/1933 [H1N1] strain) were kindly provided by Dr. Toru Takimoto at University of Rochester School of Medicine and Dentistry. Vesicular Stomatitis Virus (VSV) was purchased (Cat # VR-3340, ATCC). When cells reached >80% confluence, Influenza A viral stock (MOI = 1) was added to cell media and incubated with cells for 36 or 48 hrs respectively. SeV and VSV viral stock (MOI = 1) was incubated with cells for 24 hrs.

### Cell transfection

Poly(I:C) was transfected with the Fugene-6 transfection reagent (Cat. # E2691, Promega). Cells were incubated with Poly(I:C) for 24 hrs. For gene knockdown, siRNA (10 nM) was reversely transfected in cells using Lipofectamine™ RNAiMAX transfection reagent (Cat. # 13778030, Invitrogen) for 72 hrs. The following siRNAs were used: siNAT10-1 (Cat. # S30491, Life Technologies); siNAT10-2 (Cat. # S30492, Life Technologies). For the negative control, we used the Silencer™ Negative Control No. 4 siNT (Cat. # AM4641, Invitrogen).

### Luciferase assay

HEK293T cells were transfected with siRNAs. At 72 hrs post-transfection, cells were transfected with plasmids including pGL3-IFN-β-luciferase reporter vector (Cat. # 102597, Addgene) and pRL-TK renilla luciferase vector (Cat. # E2241, Promega) along with pTRIP-SFFV-mtagBFP-2A RIGI WT (Cat. # 167289, Addgene) using the TurboFect™ transfection reagent (Thermo Scientific) for 24 hrs. Cells were treated with Poly(I:C) for another 24 hrs. The Dual-Glo® luciferase system (Cat. # E2920, Promega) was used for quantification of firefly and renilla luminescence. Luminescence was measured by the Cytation 5 multimode reader (BioTek). Relative luciferase unit (RLU) was calculated by normalizing firefly by renilla signal.

### Protein immunoblotting

Cells were lysed in the RIPA buffer (Cat. #20-188, Millipore) containing protease inhibitor cocktail (Cat. # A32965, Thermo Scientific) on ice. With brief sonication, the total protein amount in cell lysate was measured using BCA assay kit (Cat. #23225, Thermo Scientific) for the loading normalization. Lysates were mixed with the SDS loading buffer containing 5% β-mercaptoethanol (Cat. #60-24-2, Acros Organics) and boiled. Protein samples were loaded on the Novex™ WedgeWell™ 4-20% SDS-PAGE Tris-Glycine gel (Invitrogen) or the Mini-Protean TGX gel (Biorad), and transferred to the iBlot™ 2 Transfer Stacks PVDF (Invitrogen) or the RTA transfer kit PVDF (Biorad) using the iBlot 2 Dry Blotting System (Cat. # IB21001, Thermo Scientific) or the Trans-blot Turbo (Cat. # 1704150, Biorad). Membranes were blocked with 5% BSA in PBS for 1 hr and incubated with the primary antibodies at 4°C for overnight, followed by the incubation with HRP-conjugated secondary antibodies for 1 hr. The membranes were washed three times with PBST (PBS with 0.5% Tween 20). The membranes were developed using the Clarity Max ECL substrate (Cat. # 1705062, Bio-Rad).

### Enzyme-linked immunosorbent assay (ELISA)

Human IFN-β DuoSet ELISA (Cat. # DY814-05, R&D Systems) was used according to the manufacturer’s protocols. Briefly, supernatants were collected from cultured cells. ELISA plates were coated with capture antibody for overnight, followed by the incubation with blocking buffer for 1 hr, supernatant for 2 hrs, detection antibody for 2 hrs, Streptavidin-HRP for 20 mins, and stopping solution for 10 mins. Between each incubation, ELISA plates were washed 4 times with the washing buffers. Absorbance at 450 nm was quantified using the Cytation 5 multimode reader. IFN-β concentrations were calculated based on a standard curve.

### Cell viability assay

Cell viability was measured using ATP-based CellTiter-Glo® (Cat. # G7571, Promega) by following manufacturer’s instructions. Briefly, Cell-Titer-Glo was added to each cell in media at 1:1 ratio in the opaque 96-well plate. The plate was incubated for 10 mins on a rocking shaker protected from the light. The luminescent signal was analyzed by the Cytation 5 reader.

### Chromatin immunoprecipitation (ChIP) assay

Cells were cross-linked with 0.5% paraformaldehyde, and the reaction was quenched using 100 mM glycine. Cell lysis was performed on ice for 10 mins in CE buffer (10 mM HEPES-KOH, 60 mM KCl, 1 mM EDTA, 0.5% NP-40, 1 mM DTT, pH 7.9) supplemented with a protease inhibitor cocktail. Nuclei were pelleted in SDS lysis buffer (1% SDS, 10 mM EDTA, 50 mM Tris-HCl, pH 8.1) containing protease inhibitors by centrifugation at 700 × g for 10 mins at 4°C. Nuclear lysates in SDS buffer were subjected to sonication for 2 mins to fragment genomic DNAs and diluted with the ChIP dilution buffer (0.01% SDS, 1% Triton X-100, 1.2 mM EDTA, 16.7 mM Tris-HCl, 150 mM NaCl, pH 8.1) containing protease inhibitors. The lysates were incubated overnight at 4°C with specific antibodies or control rabbit IgG (Cat. # sc-2025, Santa Cruz). Protein A/G beads (Cat. # 88803, Pierce) were pre-blocked with 0.5 mg/ml BSA and 0.125 mg/ml herring sperm DNA (Cat. 15634-017, Invitrogen) at room temperature for 1 hr and then added to the mixture for incubation for another 2 hrs at 4°C. The beads were washed sequentially with the following buffers: low salt wash buffer (0.1% SDS, 1% Triton X-100, 2 mM EDTA, 20 mM Tris-HCl, 150 mM NaCl, pH 8.1); high-salt wash buffer (0.1% SDS, 1% Triton X-100, 2 mM EDTA, 20 mM Tris-HCl, 500 mM NaCl, pH 8.1); LiCl buffer (0.25 M LiCl, 1% NP-40, 1% Na-deoxycholate, 1 mM EDTA, 10 mM Tris-HCl, pH 8.1); and TE buffer (10 mM Tris-HCl, 0.1 mM EDTA, pH 8.1). Protein samples were eluted using the fresh elution buffer (1% SDS, 0.1 M NaHCO3) at room temperature. The eluates were incubated at 65°C overnight with 0.2 M NaCl to dissociate protein-DNA complexes, followed by treatment with proteinase K (Cat. # EO0491, Thermo Fisher Scientific) at 50°C for 2 hrs for protein digestion. DNAs were extracted using UltraPure™ Phenol:Chloroform:Isoamyl Alcohol (Cat. # 15593031, Thermo Fisher Scientific) by following manufacturer’s instructions, and the resulting DNA pellets were resuspended in water for subsequent qPCR analysis.

### Quantitative PCR (qPCR)

Total RNAs were extracted from cells using the NucleoSpin RNA extraction kit (Cat. # 740955.250, MACHEREY-NAGEL). Subsequently, 500 ng of RNAs were reverse transcribed into cDNAs using the iScript™ cDNA synthesis kit (Cat. # 1708890, Bio-Rad). Real-time qPCR was carried out using the iTaq™ Universal SYBR® Green Supermix (Cat. # 1727125, Bio-Rad) on a Bio-Rad CFX Connect qPCR system. The thermal cycling conditions were as follows: an initial denaturation step at 95°C for 10 mins, followed by 50 cycles of 95°C for 15 secs and 60°C for 1 min. The relative cDNA level of the gene was normalized to GAPDH using the 2^-ΔΔCt method: 2 ^(ΔCT of targeted gene - ΔCT of GAPDH)^. Values were normalized to the control using the same method: 2^(ΔCT of targeted gene - ΔCT of IgG pull-down)^. The qPCR primers were listed (**Table S2**).

### RNA sequencing (RNA-seq) analysis

Total RNAs were extracted from cells using the NucleoSpin RNA extraction kit. RNA samples were submitted to GENEWIZ (South Plainfield, NJ) for quality control and further procedures. Briefly, RNA samples were processed with rRNA removal via poly-A selection, followed by library preparation. Sequencing was performed on an Illumina HiSeq platform using a 2 × 150 bp paired-end configuration, achieving more than 20 million reads per cell sample. Differential gene expression analysis was conducted using the DESeq2 package. Heatmaps and pathway analysis were generated with the R package *pheatmap*, and the *clusterProfiler* package.

### Immunofluorescence assay (IFA)

Cells were fixed with 4% paraformaldehyde for 10 mins at room temperature and permeabilized with 0.2% Triton X-100 for 15 mins. Blocking was performed with D-PBS containing 5% FBS for 1 hr. Cells were incubated overnight at 4°C with the primary antibodies in D-PBS containing 2.5% FBS, followed by incubation for 1 hr at RT with Alexa Fluor 488-conjugated (Cat. #A11008, invitrogen) or Alexa Fluor 568-conjugated (Cat. #A11004, invitrogen) secondary antibodies. Nuclei were stained with Hoechst 33342 (Thermo Scientific) for 10 mins at room temperature. Imaging was carried out by using a confocal microscope, and the percentage of virus-infected cells was quantified by using ImageJ.

### Statistics

Statistical analysis was performed by using the GraphPad PRISM. Data were presented as Mean ± standard error of the mean (S.E.M.) of biological or technical repeats from at least 3 independent experiments. * p<0.05, ** p<0.01, *** p<0.001, or **** p<0.001 indicated the significant difference analyzed by unpaired two-tailed t-test, one-way or two-way ANOVA and Bonferroni multiple comparison test.

## Acknowledgments

We thank Dr. Toru Takimoto (University of Rochester School of Medicine and Dentistry) for providing Influenza A virus (A/WSN/1933 [H1N1] strain) and SeV. We also thank Dr. Jacob Yount, Dr. Ruben C Petreaca, and Dr. David Bisaro from Ohio State University for helpful comments. This study was funded by NIH research grants R01DA059538, R01MH134402, and R56AI181631 to JZ, and R01CA260690 to NS.

## Author contributions

JZ, NS, and YP conceived and designed this study; YP performed most of the experiments; JH and ZW provided experimental support; YP, NS, and JZ analyzed the results; JH, ZW, GF, DZ, ZH, KS, TL, JC, and YL, contributed reagents and/or provided advice regarding this study; YP, NS, and JZ wrote the manuscript; NS and JZ reviewed the manuscript; and JZ and NS supervised the entire study.

## Data and materials availability

All data needed to evaluate the conclusions in the paper are present in the paper and/or the Supplementary Materials. Additional data and materials related to this paper may be requested from the authors.

