## Supplementary material for "NAT10 Suppresses RNA Sensing Induced IFN-β Transactivation to Promote Viral Infection via Interfering with IRF3 Activities": Suppl Figures

### Figure S1

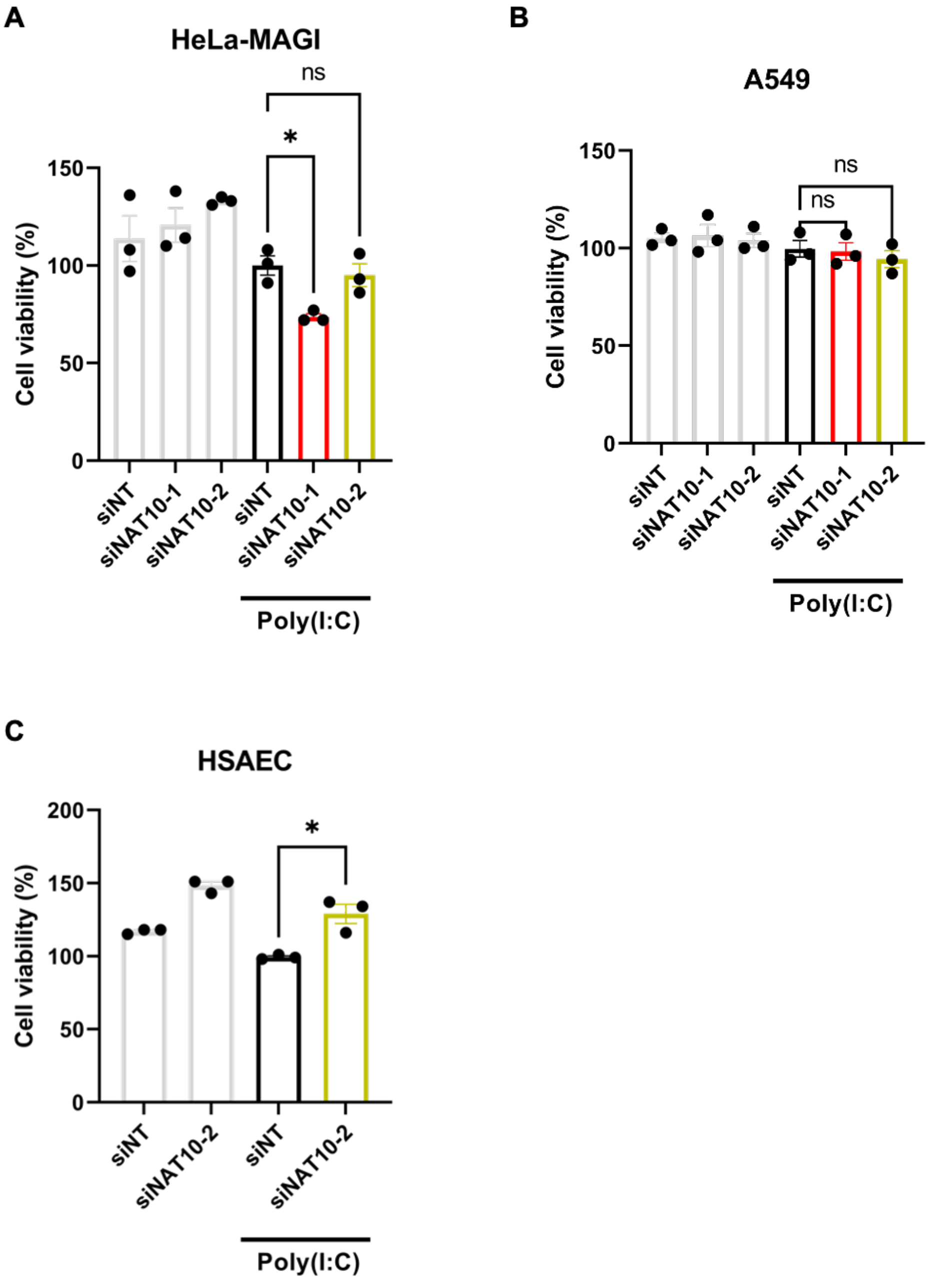


**Figure S1.** (**A-C**) Viability of HeLa-MAGI (A), A549 (B), and HSAEC (C) cells transfected with indicated siRNAs at day 3 without or with poly(I:C) treatment. The results were presented as the Mean ± S.E.M.; ns, not significant. *$P < 0.05$, ANOVA and Bonferroni multiple comparison test.

**Figure S2**

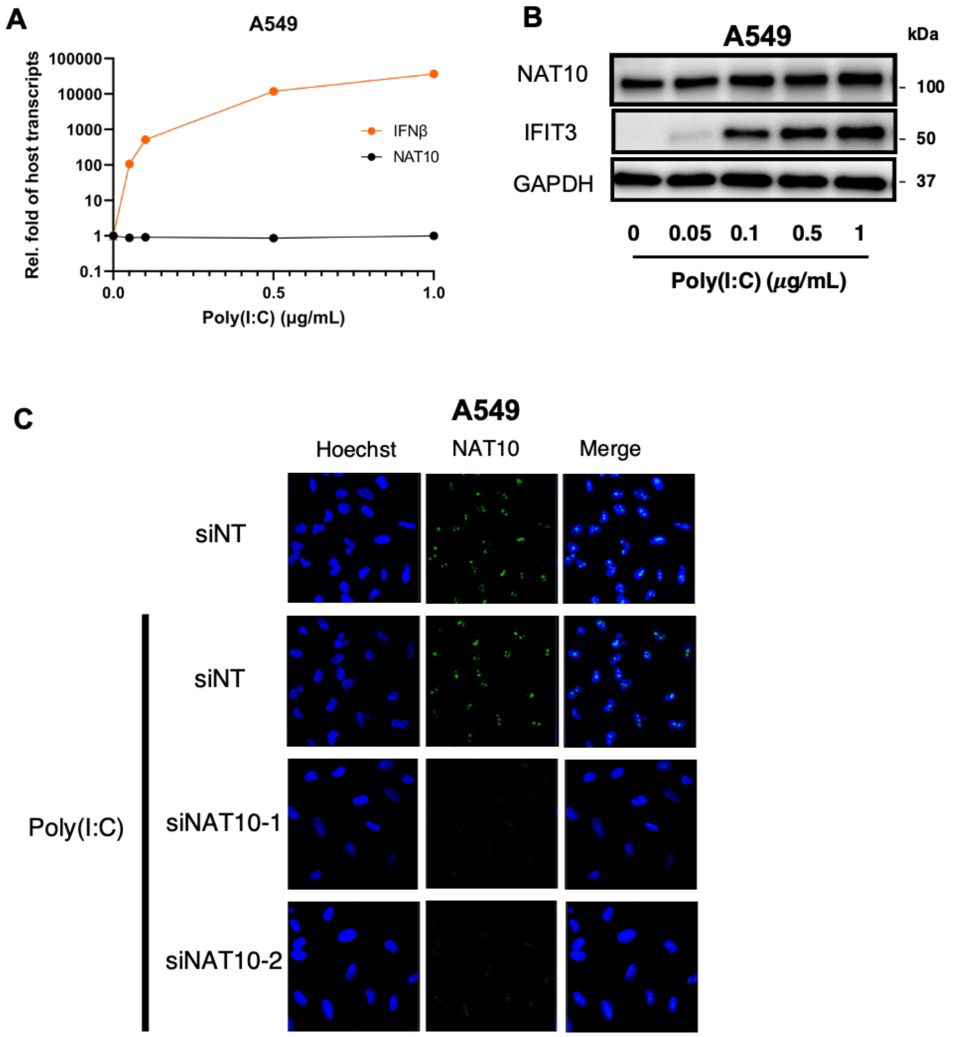


**Figure S2.** (**A**) A549 cells were treated with poly(I:C) at a series of doses (0, 0.05, 0.1, 0.5, 1μg/mL). Expression of NAT10 or IFN-β was measured by RT-qPCR. (**B**) Cell samples in (A) were analyzed by immunoblotting to quantify NAT10 and IFIT3 proteins. GAPDH was used as a loading control. (**C**) Protein expression of NAT10 was visualized by immunofluorescence (Alexa 488) in A549 cells transfected with indicated siRNAs and treated with poly(I:C).

**Figure S3**

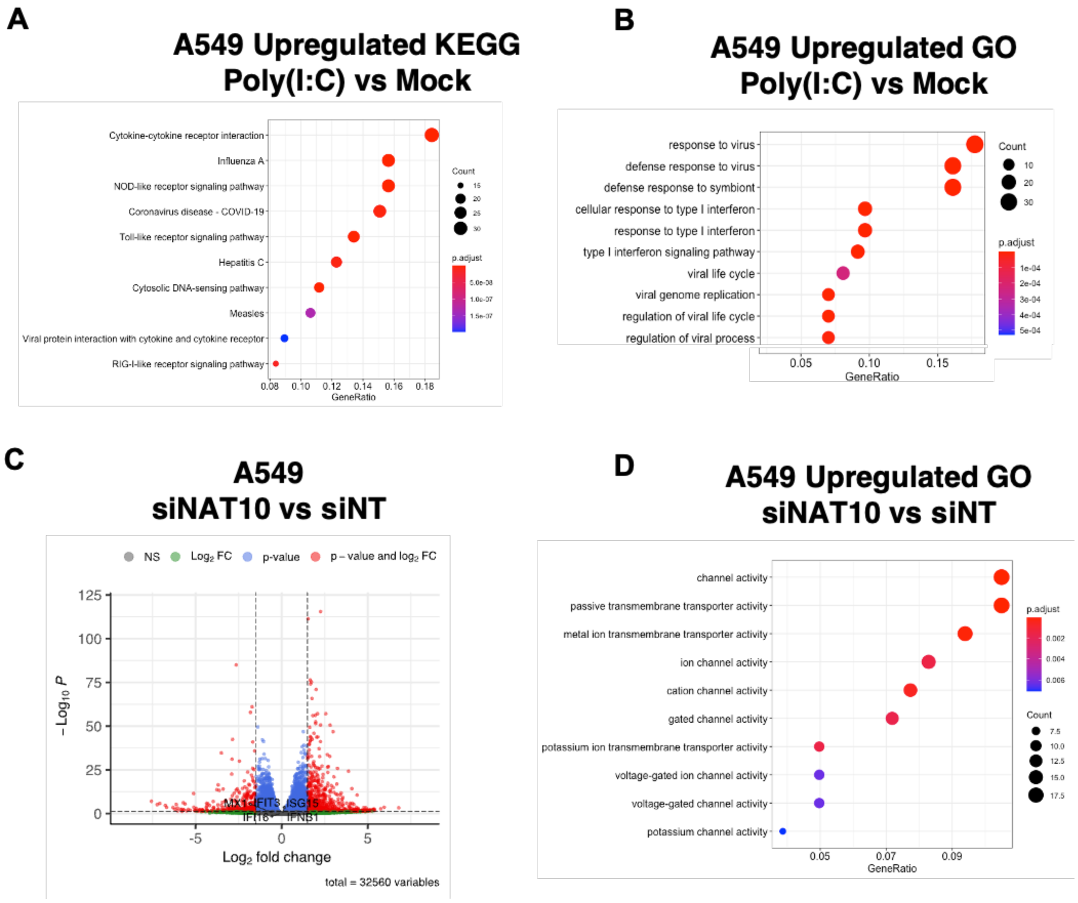


**Figure S3.** (**A, B**) KEGG (A) and Go (B) analysis of DEGs induced by Poly(I:C) treatment in A549 cells. **(C, D)** Volcanoplot (C) and GO analysis (D) of upregulated DEGs due to NAT10 depletion by its siRNAs in A549 cells without poly(I:C) treatment.

### Figure S4

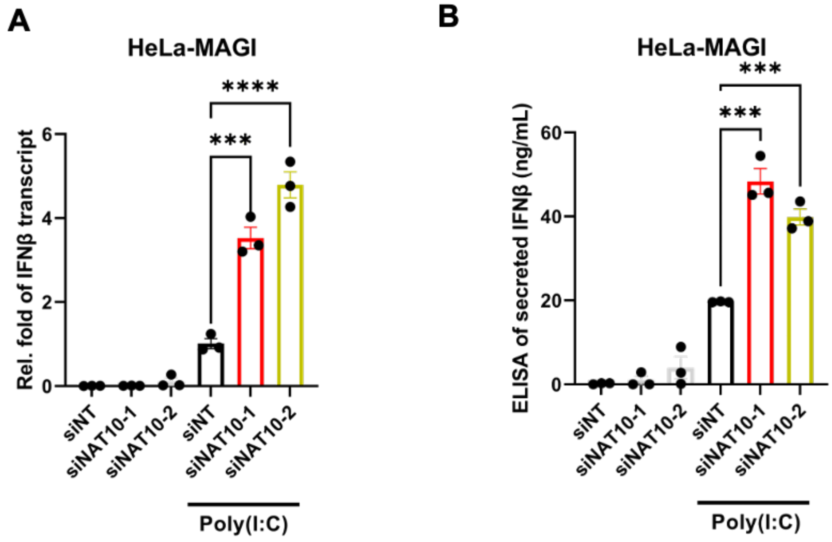


**Figure S4. (A)** HeLa-MAGI cells were transfected with indicated siRNAs, followed by the poly(I:C) treatment (1μg/mL). Expression of IFN-β was measured by RT-qPCR. (**B**) Cell supernatants in (A) were collected to quantify the secreted IFN-β protein by ELISA. The results were presented as the Mean ± S.E.M.; ns, not significant. ***$P < 0.001$, ****$P < 0.0001$, ANOVA and Bonferroni multiple comparison test for (A,B).

**Figure S5**

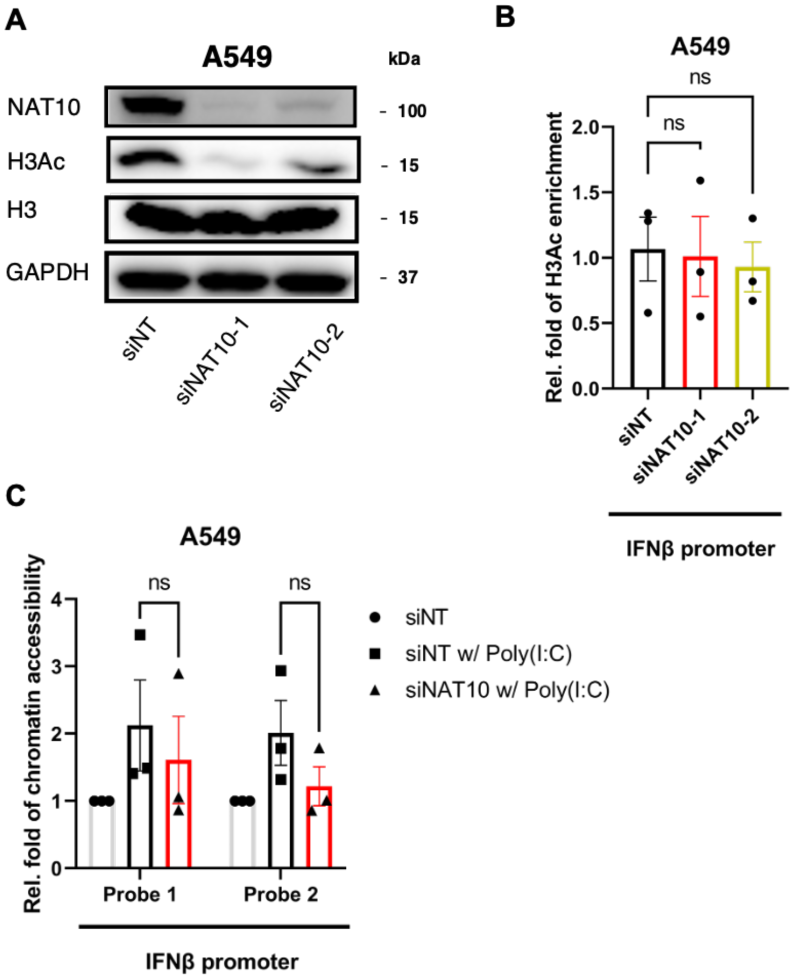


**Figure S5.** (**A**) A549 cells were transfected with indicated siRNAs, followed by the poly(I:C) treatment (1μg/mL). Protein level of NAT10, histone H3, acetylated H3 (H3Ac) was quantified by immunoblotting. GAPDH was used as a loading control. (**B**) Cell samples in (A) were processed for ChIP-PCR to determine the H3Ac association at the promoter region of IFN-β. (**C**) A549 cells were transfected with NAT10 or non-targeting siRNAs (1:1 mixture of siNAT10-1 and siNAT10-2, siNT), followed by poly(I:C) treatment. Cell samples were processed for measurement of chromatin accessibility at the promoter region of IFN-β. The results were presented as the Mean ± S.E.M.; ns, not significant., two-tailed Student t-test for (B,C).

### Figure S6

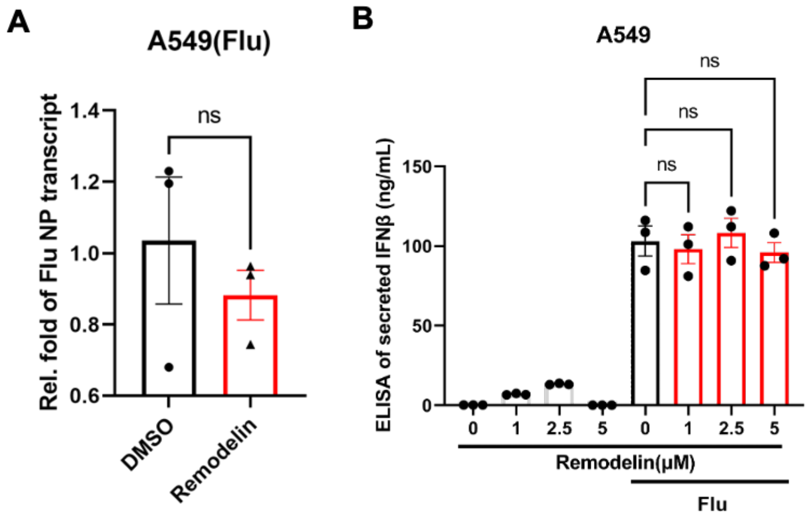


**Figure S6.** (**A**) A549 cells were treated with remodelin (5 uM), followed by challenge with influenza viruses. Expression of flu NP was measured by RT-qPCR. (**B**) Supernatants from A549 cells were treated with remodelin at indicated doses, either un-infected or infected with influenza, which were collected to quantify the secreted IFN-β protein by ELISA. The results were presented as the Mean ± S.E.M.; ns, not significant, ANOVA and Bonferroni multiple comparison test for (B) or two-tailed Student t-test for (A).

**Tables S1.** Differentially expressed genes (DEGs) induced by NAT10 depletion in A549 cells treated with poly(I:C).

**Tables S2.** Sequences of qPCR primers used in this study.
